# Global diversity of enterococci and description of 18 novel species

**DOI:** 10.1101/2023.05.18.540996

**Authors:** Julia A. Schwartzman, Francois Lebreton, Rauf Salamzade, Melissa J. Martin, Katharina Schaufler, Aysun Urhan, Thomas Abeel, Ilana L.B.C Camargo, Bruna F. Sgardioli, Janira Prichula, Ana Paula Guedes Frazzon, Daria Van Tyne, Gregg Treinish, Charles J. Innis, Jaap A. Wagenaar, Ryan M. Whipple, Abigail L. Manson, Ashlee M. Earl, Michael S. Gilmore

## Abstract

Enterococci are commensal gut microbes of most land animals. They diversified over hundreds of millions of years adapting to evolving hosts and host diets. Of over 60 known enterococcal species, *Enterococcus faecalis* and *E. faecium* uniquely emerged in the antibiotic era among leading causes of multidrug resistant hospital-associated infection. The basis for the association of particular enterococcal species with a host is largely unknown. To begin deciphering enterococcal species traits that drive host association, and to assess the pool of *Enterococcus*-adapted genes from which known facile gene exchangers such as *E. faecalis* and *E. faecium* may draw, we collected 886 enterococcal strains from nearly 1,000 specimens representing widely diverse hosts, ecologies and geographies. This provided data on the global occurrence and host associations of known species, identifying 18 new species in the process expanding genus diversity by >25%. The novel species harbor diverse genes associated with toxins, detoxification, and resource acquisition. *E. faecalis* and *E. faecium* were isolated from a wide diversity of hosts highlighting their generalist properties, whereas most other species exhibited more restricted distributions indicative of specialized host associations. The expanded species diversity permitted the *Enterococcus* genus phylogeny to be viewed with unprecedented resolution, allowing features to be identified that distinguish its four deeply rooted clades as well as genes associated with range expansion, such as B-vitamin biosynthesis and flagellar motility. Collectively, this work provides an unprecedentedly broad and deep view of the genus *Enterococcus*, potential threats to human health, and new insights into its evolution.

**SIGNIFICANCE:** Enterococci, host-associated microbes that are now leading drug-resistant hospital pathogens, arose as animals colonized land over 400 million years ago. To globally assess the diversity of enterococci now associated with land animals, we collected 886 enterococcal specimens from a wide range of geographies and ecologies, ranging from urban environments to remote areas generally inaccessible to humans. Species determination and genome analysis revealed host associations from generalists to specialists, and identified 18 new species, increasing the genus by over 25%. This added diversity provided greater resolution of the genus clade structure, identifying new features associated with species radiations. Moreover, the high rate of new species discovery shows that tremendous genetic diversity in Enterococcus remains to be discovered.

## INTRODUCTION

Enterococci are an unusually rugged and environmentally persistent genus (Gaca Anthony O. and Lemos José A., 2019). This characteristic appears to have been of selective advantage as animals emerged from the sea and colonized land, and now contributes to the spread of enterococci as leading causes of hospital-associated infection (Lebreton et al., 2017). Enterococci are among the most widely distributed members of gut microbiomes in land animals, from invertebrates to humans (Lebreton et al., 2014). Their occurrence in animals with widely varying gut physiologies, diets and social habits provides a unique opportunity to explore how diverse host backgrounds drive microbiome composition.

Pioneering surveys in the 1960’s and 1970’s by Mundt and colleagues provided early evidence for the widespread occurrence of enterococci in diverse hosts including mammals and birds (Mundt, 1963a), insects (Martin and Mundt, 1972), and animal-inhabited environments (Mundt, 1961; Mundt et al., 1958). However, at the time of those studies, the genus *Enterococcus* had yet to be recognized as distinct from *Streptococcus* (Schleifer and Kilpper-Balz, 1984) and was resolved into species at low resolution by a small number of metabolic tests (Facklam, 1973). Although evidence of diversity among the enterococci was found, the inability to precisely assess strain differences limited the ability to associate well defined *Enterococcus* species with particular hosts. Genomics now provides a high-resolution tool capable of detecting differing traits between species of microbes from varying hosts, and for quantifying the extent of their divergence.

The goal of the current study was to sample the Earth broadly for enterococci from diverse hosts, geographies, and environments, to compare the content and degree of divergence of their genomes, toward the broader goal of understanding the mechanisms that drive association with particular hosts. This was also done to gain insights into the scope and magnitude of the *Enterococcus* gene pool on a planetary scale. To achieve these goals, enterococci from 381 unprocessed animal samples, as well as 456 enterococci isolated by contributors from diverse sources, were collected and taxonomically identified at the DNA sequence level. The entire genomes of strains exhibiting sequence diversity suggestive of distant relationship to any known species, were then sequenced in their entirety. This identified 18 new species of *Enterococcus* and 1 new species of the ancestrally related genus *Vagococcus*. Genome sequence analysis also showed that substantial enterococcal diversity remains to be discovered, most prominently in arthropod hosts and insectivores. Further, genetic novelty was found not only in novel species, but also circulating in well-known *Enterococcus* species, including a novel BoNT-type toxin (Zhang et al., 2018) and a new family of pore-forming toxins (Xiong et al., 2022), highlighting the importance of broader knowledge of the enterococcal gene pool.

## RESULTS

### Broad survey samples *Enterococcus* host diversity

To understand the breadth of enterococcal diversity, we examined little-sampled (non-clinical, non-human) environments, including those minimally impacted by human habitation or pollution. To maximize global coverage, we assembled the Enterococcal Diversity Consortium (EDC), an international group of scientists and adventurers, who contributed 456 colony purified presumptive enterococci and 579 whole specimens, typically insects and scat, in addition to commercially procured samples (Table S1). The collection included isolates from a wide range of hosts and host diets (e.g., carnivores versus herbivores), geographies and environments (e.g., captive versus wild). The diversity of sources spanned penguins migrating through sub-Antarctic waters (Prichula et al., 2020, 2019), duiker and elephants from Uganda; insects, bivalves, sea turtles, and wild turkeys from Brazil to the United States; kestrel and vultures from Mongolia; wallaby, swans, and wombats from Australia; and zoo animals and wild birds from Europe.

From the 579 whole specimens, both CHROMagar Orientation agar (Merlino et al., 1996) and bile-esculin azide agar (Facklam, 1973) were used to culture presumptive enterococci in order to minimize potential selection bias against natural enterococcal isolates with unknown properties. To recover enterococci potentially present in low abundance, we performed isolations both directly from samples and from enrichment cultures. Some specimens yielded multiple colony morphologies, and in those cases each morphotype was separately analyzed (Table S1). At least one presumptive enterococcal isolate was culturable from 55% of samples (318 of 579), including extracts from dead insects, wild animal feces, tissue swabs, or samples of water and soil likely contaminated with animal fecal matter. These 318 presumptive *Enterococcus*-positive samples yielded 430 morphologically distinct colony types. Together with the 456 presumptive enterococcal isolates contributed as pure cultures by EDC members, this resulted in an analytical set of 886 presumptive enterococci (Table S1, Tab 1) derived from 774 different sample sources.

### Preliminary screen for species diversity

Species-level diversity within the genus *Enterococcus* is not well resolved by nucleotide polymorphisms in the 16S rRNA gene (Devriese et al., 2002). Thus, as an initial measure of taxonomic diversity, we developed a high-resolution PCR amplification and amplicon sequencing protocol. An internal 97 bp polymorphic fragment of an RNA methylase gene (Fig. S1A), designated EF1984 in the *E. faecalis* V583 genome and previously found to be core to the *Enterococcus* genus (Lebreton et al., 2017), flanked by short conserved sequence stretches that could be used for amplification, was selected for initial assessment of strain diversity. Sequence variation within this 97 bp diversity locus (DL) proved better able to discriminate species than variability within the entire 16S rRNA gene (Fig. S1B). Further, sequence polymorphism within the DL recapitulated the phylogenetic relationships between the 47 known enterococcal species based on genome-wide average nucleotide identity (ANI) (Fig. S1C).

### DL variation presumptively identified novel species

Colonies of all 886 presumptive enterococcal isolates were subjected to DL amplification and sequencing. Of those, 34 isolates yielded no product. Amplification and sequencing of the full-length 16S rRNA gene of those 34 showed them to be *Carnobacterium* (9/34), *Lactobacillus* (10/34), or *Vagococcus* (14/34), all closely related to *Enterococcus*, likely accounting for their growth on selective media used.

DL positive enterococci (852 of 886 isolates) derived from 41 taxonomic orders of animal hosts from 16 countries on 6 continents, representing many climatic zones (Fig. 1A). Largely reflecting representation in the collection, positively identified enterococcal isolates were obtained from mammals (29%), birds (28%), insects (18%), reptiles, and amphibians (9%), coastal fish (4%), bivalves (2%), and gastropod mollusks (2%) (Fig. 1B). Samples derived from primary consumers (e.g., herbivores) as well as predators and scavengers (Fig. 1C). Over half of the isolates (53%) derived from wild environments with very low human activity (Fig. 1D).

**Figure 1.**
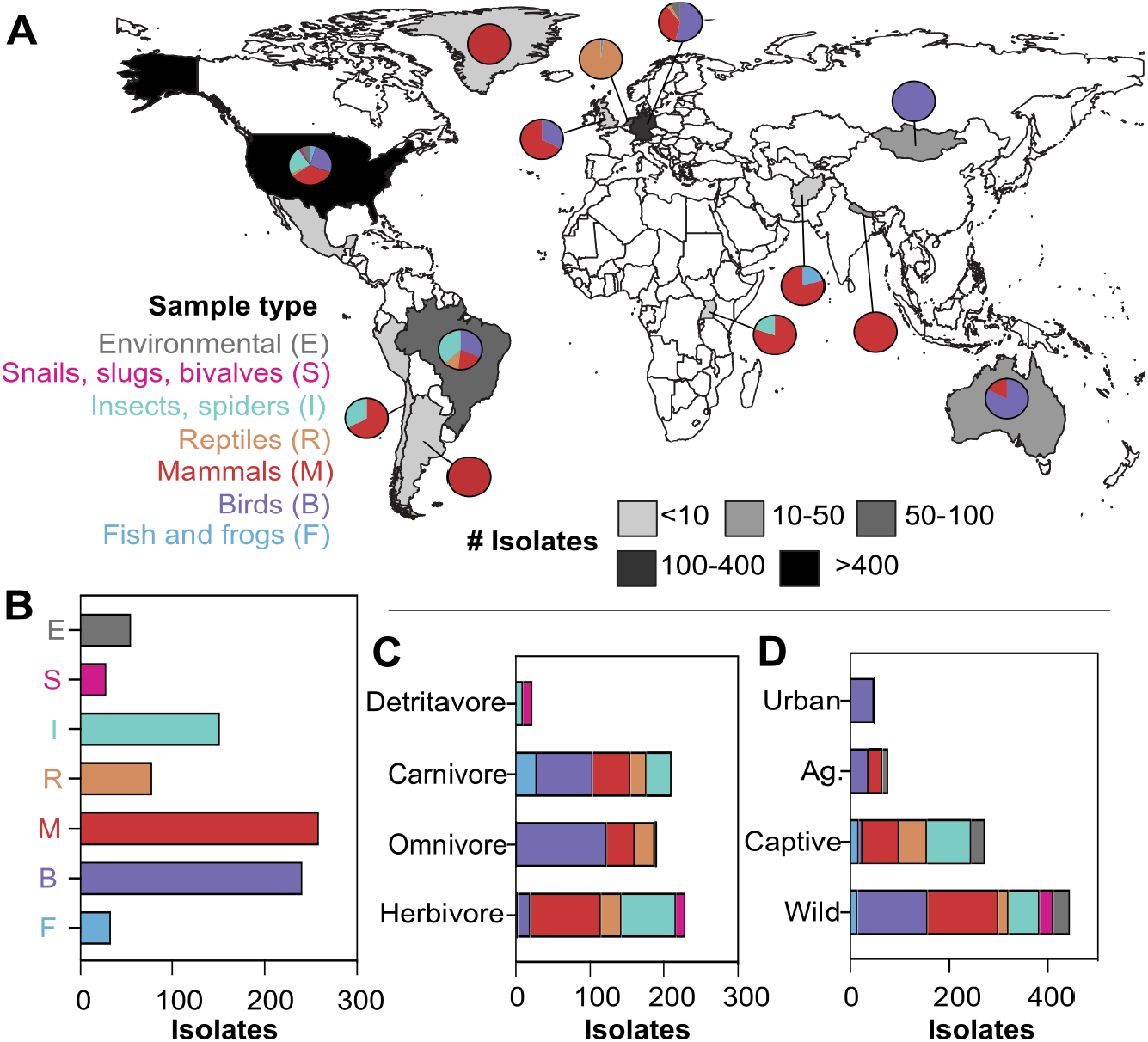
Sources for 852 *Enterococcus* isolates. **A**) Isolates derived from 16 countries, with the largest numbers collected in the United States, Germany, the Netherlands, and Brazil as reflected by grey shading. Pie chart insets show the host taxonomic distribution, colored as indicated in the key. **B-D**) Distribution of host-derived samples grouped by taxonomy, diet, and environment. Colors are the same as in panel A. **B**) Enterococcal isolates were obtained from environmental sources, as well as from eleven classes of animals representing three phyla: mollusks (snails, slugs, bivalves), arthropods (insects, millipedes, spiders), and vertebrates (reptiles, mammals, birds, fish, and frogs). The common names of major classes are shown; metadata are listed in Table S1. **C**) Enteric samples and feces were derived from animals with diverse diets. Where known, the diet of the animal was classified based on trophic role: detritivore, carnivore, omnivore, or herbivore. **D**) Samples derived from locations differing in human impact. Habitats were defined as: wild, no human activity; captive, animals housed in enclosures with human contact excluding agricultural animals; agricultural animals housed in enclosures and bred by humans for agricultural purposes (Ag); urban, wild animals living in cities or towns.

Strains belonging to the same known species all shared DL sequence variations of 4 bp or less, and DL sequence variation was able to resolve known fine scale differences between *E. faecium* clades A and B (known to share ∼94% ANI, (Belloso Daza et al., 2021; Lebreton et al., 2013). Most DL sequences (96%, 824/853) matched with fewer than 4 SNPs to one of 65 previously identified enterococcal species for which a genome had been sequenced (Fig. 2A; Table S1), to which these closely related isolates were presumptively assigned. The most frequently encountered species in our collection were: *E. faecalis* (340/853; 40%), *E. faecium* (125/853; 15% of isolates, including 66 [9%] clade A and 59 [7%] clade B), *E. mundtii* (119/853; 13%), *E. casseliflavus* (81/853;10%), and *E. hirae* (68/853; 8%). Importantly, this approach identified 27 isolates possessing 19 different DL sequences that exceeded the threshold for likely membership in a known species (>4 SNPs from any known species), identifying them as potentially novel (Fig. 2B; Table S1). These 27 isolates derived from insects (14 isolates), birds (9 isolates), and herbivorous reptiles (4 isolates) (Fig. 2C). In contrast, isolates of the most frequently encountered species - *E. faecalis, E. faecium, and E. mundtii* - derived mainly from mammals and birds, but were not exclusive to those hosts (Fig. 2C).

**Figure 2.**
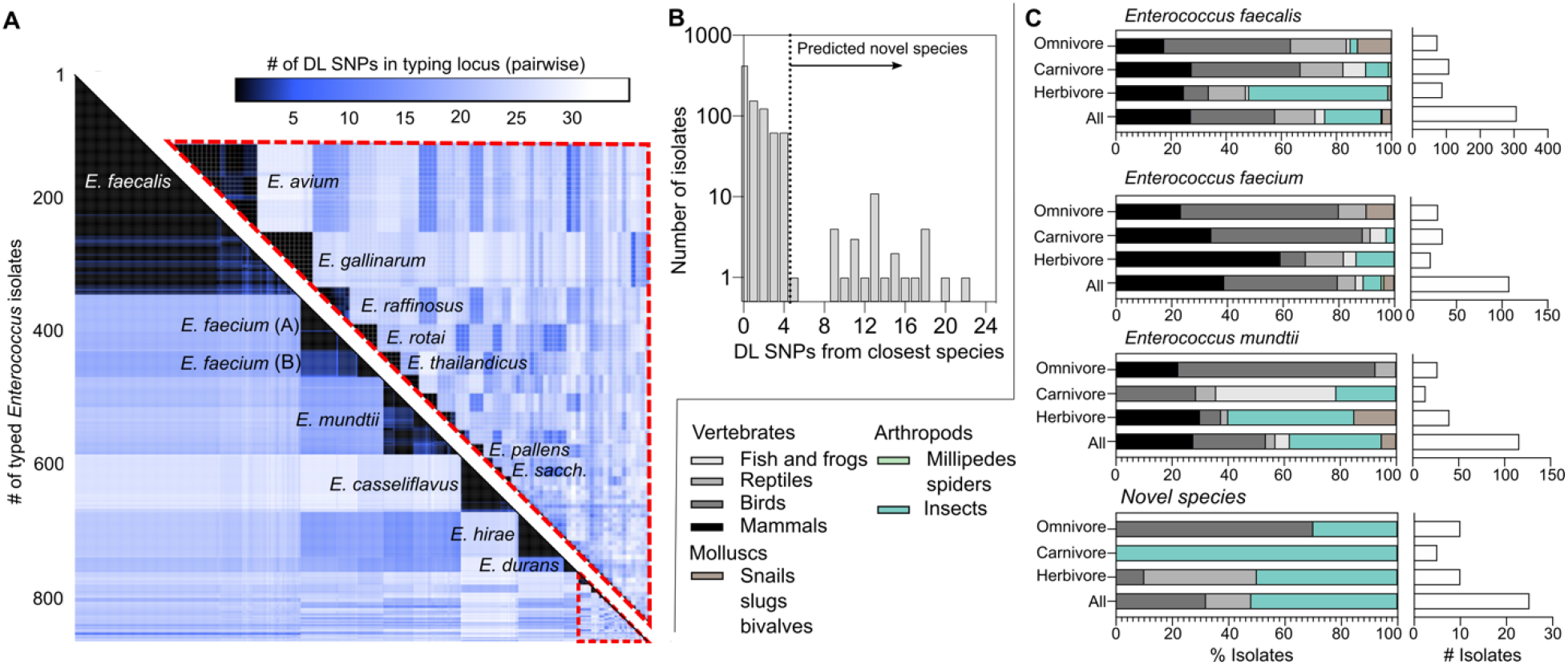
Diversity of *Enterococcus* species isolated from host classes. **A**) Heat map displaying the pairwise relatedness of the 852 putative *Enterococcus* isolates characterized by this study. Color represents the number of single-nucleotide polymorphisms (SNPs) in the DL locus between pairs of isolates. Red dashed enclosed triangle highlights the identity of rarely encountered isolates expanded in the inset. **B**) Histogram of isolates binned by the number of SNPs in the DL locus, compared to the closest known species. Vertical dashed line is placed greater than 4 SNPs, the threshold chosen to prioritize genome sequencing of diverse isolates and initially predict novel species. **C**) Host derivation of isolates, highlighting isolation sources for the three most abundant species sampled in this collection, as well as for diverse novel species.

### Species boundaries and placement within the clade structure of *Enterococcus*

Genomes of isolates presumptively identified as diverse in the DL locus when compared to their closest relative (n=22), as well as 16 *Enterococcus* and 9 *Vagococcus* isolates for which few genomes occur in public databases, were sequenced to generate high quality draft genome assemblies (Table S2). Presumptively diverse isolates that duplicated others exactly in DL sequence and derived from the same host sample type/sample location, and thus likely represented duplicate strains, were not sequenced. For each of the 47 newly sequenced *Enterococcus* and *Vagococcus* genomes, and for high quality existing assemblies of 65 known *Enterococcus* species and 4 known *Vagococcus* species, we calculated pairwise genome-wide average nucleotide identity (ANI) to determine species identities and explore species boundaries more accurately. The median % ANI shared by putative novel species and their nearest taxonomic neighbor was 83%, indicating that most novel species identified in this study were substantially distant and not closely related to known species. The number of DL SNPs correlated highly with genome-wide ANI (Figure S1C), validating DL sequence as an accurate predictor of novelty; 96% of isolates with >4 SNPs in the DL exhibited <95% ANI (the operationally defined species boundary (Jain et al., 2018)) with the genome of the closest identifiable species. Exceptions included two divergent *E. casseliflavus* genomes that differed by 5 DL SNPs from the reference but shared nominally >95% ANI (DIV0233, 95.2% ANI; 3H8_DIV0648, 95.5% ANI) with the *E. casseliflavus* reference genome. Additionally, despite differing from the reference DL by only 2 SNPs, three strains shared only 94.1% ANI with the reference *E. rotai* genome indicating that although the DL locus is conserved in this case, these also represent new species by ANI criteria. Of the *Vagococcus* genomes that were sequenced, 8/9 were confirmed to be strains of *V. fluvialis*, sharing between 97.3% and 99.9% ANI with the *V. fluvialis* reference genome (Table S2). One *Vagococcus* isolate was highly divergent from any known species, but taxonomically nested within the genus. This isolate, collected from infected tissue of a harbor porpoise (Table S2), shared only 78.5% ANI with the closest known species, *Vagococcus teuberi*, indicating that it also is a candidate novel *Vagococcus* species.

We previously observed that species within the genus *Enterococcus* clustered into four deep-branching clades (Lebreton et al., 2017). Since that observation was based on a much more limited dataset, we re-evaluated enterococcal phylogenetic structure, including placement of the newly described variants and species found here, along with publicly available reference enterococcal genomes added since our prior analysis. We further included an expanded set of outgroup genomes, representing close taxonomic relatives of enterococci, including *Vagococcus, Pilibacter*, and *Catellicoccus*, including 9 *Vagococcus* genomes sequenced in this study (Table S2, Fig. 3A).

**Figure 3.**
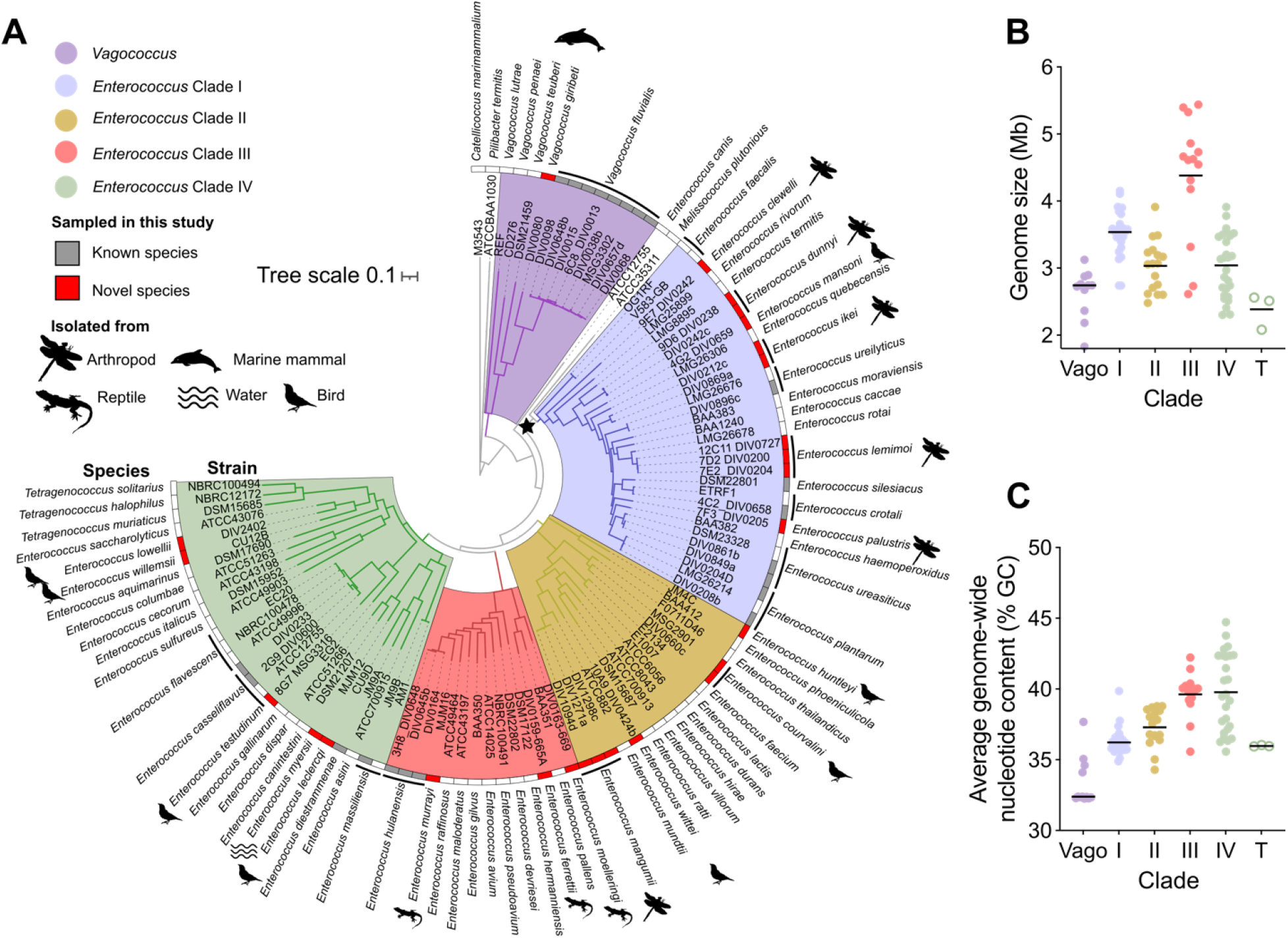
Phylogeny of known and novel Enterococcus species. **A**) Phylogenic positioning of 5 *Vagococcus* species and 65 *Enterococcus* species, including 18 novel species of *Enterococcus* and one of *Vagococcus*. Black icons show the host source from which novel species isolates were derived. Strains of the same species are indicated by black lines. Clades are denoted by colored regions, Clade I blue; Clade II yellow; Clade III red; and Clade IV green. *Vagococcus* spp. are noted by purple edges. Star indicates bootstrap support ≥ 90%. **B-C**) Distribution of genome size (B) and average %GC content (C) in our whole genome assemblies within each clade (Vago, *Vagococcus;* I-IV, *Enterococcus* Clades I-IV; T, *Tetragenococcus,* a group of divergent species clustering within *Enterococcus* Clade IV). Bars indicate the mean.

Phylogenetic placement was based on sequence conservation in 320 single-copy genes present in all genomes. This analysis showed that the closest relative of *Enterococcus* was *Vagococcus*, with *Pilibacter* and *Catellicoccus* branching earlier within the family Enterococcaceae (Fig. 3A). Despite the placement of 41 additional species in the *Enterococcus* genus (Fig. 3A), the previously observed (Lebreton et al., 2017) structure of 4 deeply branching clades of the *Enterococcus* genus was supported. Bootstrap support for this topology was strong, with the weakest being the placement of *Melissococcus plutonius* (67%) as the most ancestral branch in clade I (Fig. 3A).

The long branch length of *M. plutonius* and its small genome size (2.05Mb) reflect its unusual trajectory in becoming a highly specialized bee pathogen (Bailey and Collins, 1982). This fundamental habitat shift within a genus of largely commensal gut microbes, likely skews its positioning in the tree. The higher resolution phylogeny also places *Enterococcus canis* as an outlier, branching very early in the evolution of the genus (Fig. 3A). All 18 new enterococcal species in our collection nested within these four main radiations of *Enterococcus* species. Ten of these novel genomes were placed into the phylogeny at ancestral branch points relative to other species in the topology, meaning that we expanded the known diversity in each clade (Lebreton et al., 2017) and revealed both recent and more evolutionarily ancient diversification within clades.

### Fundamental differences between *Enterococcus* clades

With each of the 4 deep branching *Enterococcus* clades now represented by 12 or more species, we next sought to identify fundamental differences between clades to gain insight into the possible evolutionary drivers that created this phylogenetic structure. We previously noted that members of Clade III possessed unusually large genomes up to 5.4 Mb in size (*E. pallens*), whereas members of Clade IV possessed genomes less than half that size (*E. sulfureus*, 2.3 Mb) (Lebreton et al., 2017). Here we found that Clade I genomes ranged from 2.7 Mb *Enterococcus faecalis* to the 4.2 Mb *Enterococcus termitis*, with a mean size of 3.5 Mb (Fig. 3B). Clade II genomes ranged in size from 2.5 Mb *Enterococcus ratti* to 3.9 Mb *Enterococcus phoeniculicola*, with a mean of 3.0 Mb. Clade III genomes, previously thought to contain only large genomes (Lebreton et al., 2017), were revealed to range in size from the 2.6 Mb *Enterococcus hermannienesis* to *E. pallens* (5.4 Mb), with a mean of 4.4 Mb (Fig. 3B). Clade IV genomes ranged in size from the previously noted 2.3 Mb *Enterococcus sulfureus* to 3.8 Mb *E. sp. nov. testudinum* and averaged 3.0 Mb (Fig. 3B). *Tetragenococcus* genomes, which grouped within Clade IV but appear to have adapted to an ecologically distinct environment from other enterococci, were slightly smaller than the average genome within Clade IV, ranging in size from 2.1 to 2.5 Mb, with a mean size of 2.4 Mb. Interestingly, except for the anomalous pathogen, *M. plutonius*, all 4 *Enterococcus* clades exhibited a stepwise increase in mean G+C content compared to that of ancestrally related *Vagococcus* (33%): *Enterococcus* Clade I averaged 36.2% +/- 0.9%, Clade II averages 37.3% +/-1.3%, Clade III averages 39.6% +/- 1.4%, and Clade IV averages 39.8% +/- 2.7% GC (Fig. 3C).

To further characterize fundamental differences between clades, gene content was compared. The pan-genome of the *Enterococcus* genus clustered in a total of 11,086 groups of orthologous genes, excluding singletons (n= 8,017). A subset of 1,336 genes/orthogroups were shared by most (>80%) species (Table S3, Tab 1), while 417 genes present in single copy in all genomes of *Enterococcus* species composed the strict, single copy core (SCC) genome of the genus (Table S3, Tab 2). When compared to *Vagococcus*, only 5 members of the SCC were generally missing in species of that genus (EF1063, EF1150, EF2073, EF2149, EF2695; Table S3, Tab 3). Within *Enterococcus*, 67 orthogroups were enriched in Clade I and generally missing in species in Clades II to IV. Although the most commonly encountered species in this study, *E. faecalis* is clearly anomalous within Clade I as it carries genes for only 21 of the 67 orthologous groups enriched in species of that clade. Of note, all twenty-nine Clade I species, including *E. faecalis*, encoded one or more LanC-like lanthipeptide synthetase genes orthologous to *E. faecalis cylM* (EF0527), a hallmark of lanthipeptide bacteriocin biosynthetic operons that was otherwise present in only 12% of species from other clades (Fig. 4A). Further, 17 distinct clusters for the biosynthesis of lanthipeptides were uniquely found in Clade I with some (*e.g.,* clusters L3, −5, −7, −24) shared by a large fraction of Clade I species (Fig. 4A, Table S4). Only 5 lanthipeptide synthesis clusters were found in species from other clades and, in all cases, were specific to a given isolate (Table S4). Besides enrichment, Clade I was depleted of 16 orthologous groups otherwise core to the other clades of *Enterococcus*. Most are of unknown function despite forming potential functional clusters/operons (e.g., E2134_0962 – E2134_0964, Table S3).

**Figure 4.**
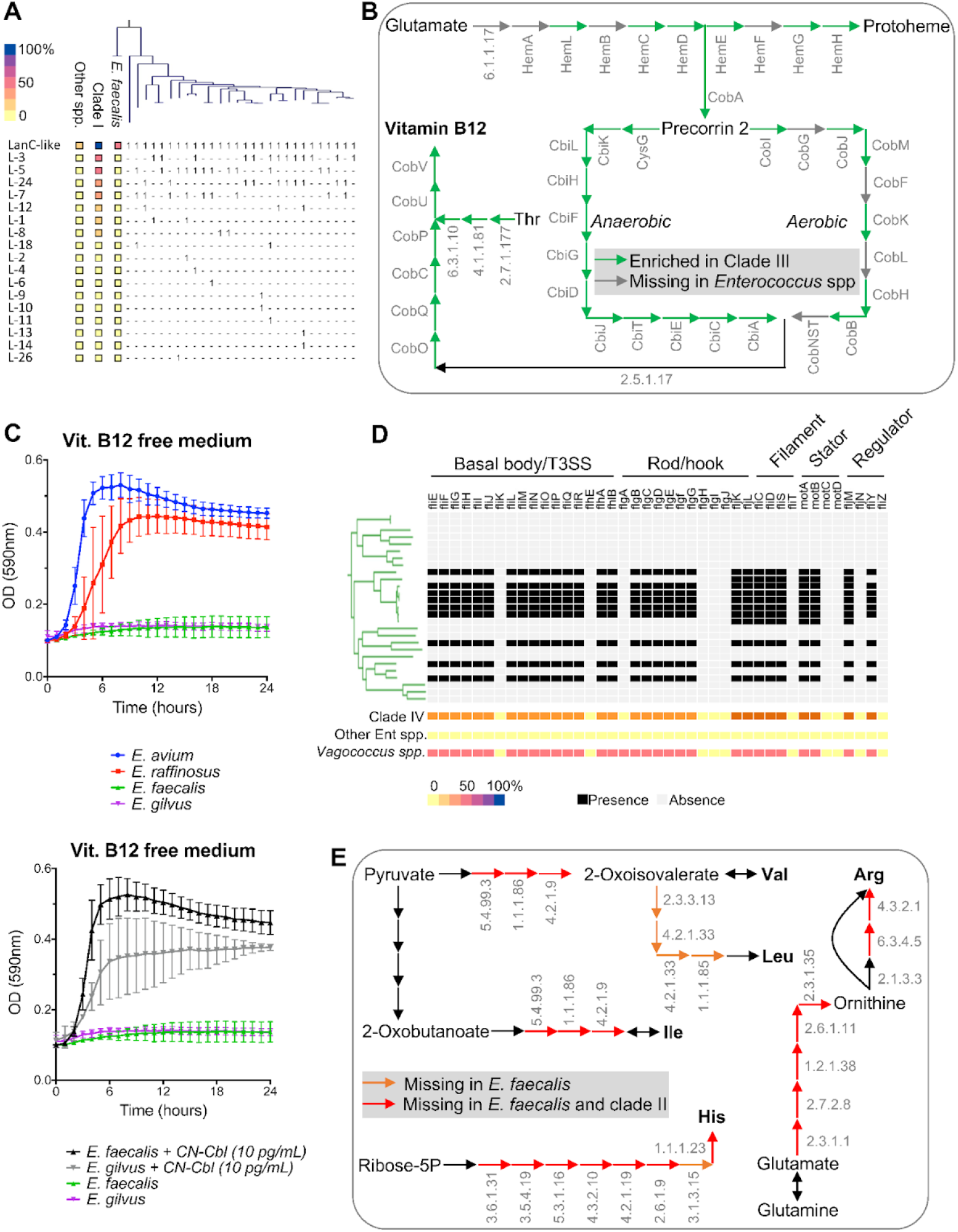
Identification of clade-specific genotypic and phenotypic traits within the *Enterococcus* genus phylogeny. **A**) Frequency of a LanC ortholog (LanC-like) as well as 17 distinct lanthipeptide synthesis clusters (Table S4) for Clade I species, non-Clade I *Enterococcus* (other spp.), and *E. faecalis.* The distribution of these orthologs and gene clusters is noted for individual members of Clade I as a heat map. The taxonomic relatedness of Clade I species is displayed as a pruned phylogenetic tree above the heatmap. The heatmap shows the relative abundance in *E. faecalis*, all Clade I species or all other *Enterococcus* spp. **B**) Enrichment of complete aerobic and anaerobic cobalamin biosynthesis pathways in a subset of Clade III species. **C**) Growth curves for a selection of isolates and species in a minimal media lacking Vitamin B12 (upper chart). For species lacking the cobalamin biosynthesis pathway, growth is rescued when 10 pg/mL of cyanocobalamin (CN-Cbl) is added exogenously (lower chart). **D**) Enrichment of flagellar biosynthesis gene clusters in subset of Clade IV species. Presence (black) and absence (grey) of specific genes is indicated for each Clade IV species (displayed as a pruned phylogenetic tree) as well as a heatmap showing their prevalence in all Clade IV species, all other *Enterococcus* spp., or a set of 12 *Vagococcus* spp. **E**) Depletion of near-complete pathways for the biosynthesis of histidine and BCAA in Clade II species together with *E. faecalis* (red arrows) or in *E. faecalis* alone (orange) compared to other *Enterococcus* spp. When available, the E.C. number of the missing enzyme is indicated.

Clade II is characterized by the smallest core genome and species within it generally lacked 42 orthologous genes relative to all other clades. Within those possessing a functional annotation, a large fraction of missing genes (∼30 %, including operon BAA382_0711 - BAA382_0714) belong to lacking amino acid biosynthesis and interconversion pathways, specifically in the biosynthesis of all three branched-chain amino acids (BCAAs), but also histidine and arginine (Fig. 4E). Of note, the majority (>80 %) of orthologous groups depleted in Clade II were also missing in *E. faecalis* despite being core to other species in Clade I, further supporting *E. faecalis* as an outlier in this phylogenetic group and highlighting a curious similarity to Clade II species. For Clades II a total of 24 orthologous groups were significantly enriched. Most are of unknown function; however, some appear to aggregate in genomically co-located clusters (e.g., E2134_1832 - E2134_1834 in *E. faecium* and most other Clade II members, Table S3).

Clade III was found to be significantly enriched in 82 orthologous groups, while being depleted in 8 ortholog groups relative to other enterococci. Most genes in both sets were annotated as of unknown function (Table S3). While not shared by the arbitrary threshold of >80% of species, a large subset (64%) of Clade III species shared genes coding for complete pathways for the aerobic and anaerobic biosynthesis of cobalamin (Fig. 4B). The presence of this metabolic pathway was noted in all species with atypically large genomes, while it is missing in the few Clade III species possessing genome sizes more typical for *Enterococcus* species. Importantly, these genes were not found outside of Clade III, and an in vitro growth experiment using a defined medium, confirmed their role in the de novo biosynthesis of the otherwise essential vitamin B12 (Fig. 4C).

Finally, species of Clade IV were enriched in only 2 orthologous gene groups of largely unknown function. However, a large subset of Clade IV species (>40%) encoded genes necessary for the biosynthesis of a flagella, which were otherwise missing in any other species throughout the genus (Fig. 4D). This cluster, first described in *E. gallinarum* and *E. casseliflavus* (Palmer, 2012), shares identity with the flagellar system in lactobacilli and a similar system (at the gene content level) was also detected in >50% of the known *Vagococcus* species (Fig. S2). Clade IV species are depleted in only 1 orthologous group compared to all other clades (Table S3).

### Functional analysis of novel species

A comparison of gene content was made between the 18 novel *Enterococcus* species discovered here and their nearest taxonomic neighbor (Table 1), to gain insight into what may have selected for speciation. Nearest neighbors were defined as the previously known species that shared the highest genome-wide ANI and sharing the shortest branch length (Fig. 3A, Table 1). Where multiple reference genomes existed, gene content was required to be present or absent in all genomes of that species. Genes were first binned into functional categories using COG annotation (Fig. S3A, Table S6). As found in our prior study (Lebreton et al., 2017), carbohydrate transport and metabolism was the largest functional category enriched in species-specific gene content (Fig. S3A). Following this, the second most enriched category was suspected defense mechanisms (8.8%), with most encoding ABC-type transporters predicted to efflux antimicrobials and other noxious compounds. Regulation of transcription (7.6%), and signaling (6.9%), largely associated with responding to sources of nutrition, lend further support to the co-evolution of enterococci along with the dietary habits of their hosts (Lebreton et al., 2017).

**Table 1.** Candidate novel *Enterococcus* species described in this study.

| Species name | Strains | Clade | Most related species | ANI | Host(s) | Novel/unique genes | Examples of unique functions present in novel genomes |
| --- | --- | --- | --- | --- | --- | --- | --- |
| *Enterococcus clewelli* | 9E7_DIV0242b | I | *E. quebecensis* | <77 | Captive African bullet cockroach | 233 | Cluster of 9 genes involved in tellurium resistance, CAMP-factor toxin |
| *Enterococcus dunnyi* | 9D6_DIV0238c; DIV0242c | I | *E. termitis* | 82.0 | Ground beetle<br>Captive African bullet cockroach | 49 | E1-E2 ATPase |
| *Enterococcus mansonii* | 4G2_DIV0659b | I | *E. quebecensis* | 82.3 | Turkey | 69 | Delta-conotoxin like protein |
| *Enterococcus ikei* | DIV0869a; DIV0212c | I | *E. ureilyticus* | 83.0 | Captive moth<br>Dragonfly | 14 | Miraculin-like protein |
| *Enterococcus lemimoi* | 12C11_DIV0722f; 7D2_DIV0200; 7E2_DIV0204 | I | *E. rotai* | 94.1 | Moth<br>Dragonfly | 20 | Glyoxalase-like domain enzymes |
| *Enterococcus palustris* | 7F3_DIV0205d | I | *E. haemoperoxidus* | 84.8 | Dragonfly | 33 | Spherulin-like cytosolic protein |
| *Enterococcus huntleyi* | JM4C | II | *E. phoeniculicola* | 80.2 | Chicken | 83 | Macrolide export-like proteins |
| *Enterococcus courvalini* | MSG2901; DIV0660c | II | *E. thailandicus* | 90.8 | Cockroach<br>Turkey | 3 | Restriction endonuclease, Asel-like |
| *Enterococcus wittei* | 10A9_DIV0425 | II | *E. mundtii* | 80.6 | Chicken | 90 | Lartarcin-like toxin, cytolysin-like toxin, Gallidermin-like protein |
| *Enterococcus mangumii* | DIV1094d; DIV1271a; DIV1298c | II | *E. mundtii* | 87.5 | Butterfly<br>Dragonfly<br>Butterfly | 38 | PAAR domain protein (possible type VI secreted effector) |
| *Enterococcus moelleringi* | DIV0163-669 | III | *E. pallens* | 80.7 | Kemp's ridley sea turtle | 131 | Acetoacetate decarboxylase and collagenase |
| *Enterococcus ferrettii* | DIV0159-665A | III | *E. pallens* | 84.6 | Kemp's ridley sea turtle | 109 | MAPKK protease toxin-like protein |
| *Enterococcus murrayi* | MJM16 | III | *E. hulanensis* | 83.7 | Kemp's ridley sea turtle | 80 | Cluster of 5 genes involved in phosphonate metabolism (pnhGHIJK) |
| *Enterococcus leclercqi* | CU9D | IV | *E. diestrammenae* | 79.2 | Chicken | 84 | Alo3-like protein (antifungal), AlwI-like restriction endonuclease |
| *Enterococcus myersii* | DIV0106b-MJM12 | IV | *E. dispar* | 83.6 | Aquarium water | 50 | Bacteriocin class II with double-glycine leader peptide |
| *Enterococcus testudinum* | 8G7_MSG3316 | IV | *E. casseliflavus* | 80.6 | Captive turtle | 69 | Putative arabinoxylan degradation, macrolide efflux, Lactococcin 972-like bacteriocin |
| *Enterococcus willemsii* | CU12B | IV | *E. saccharolyticus* | 80.1 | Chicken | 58 | Possible Beta-lactamase inhibitor gene cluster |
| *Enterococcus lowellii* | DIV2402 | IV | *E. saccharolyticus* | 85.4 | Captive cockatoo | 63 | Imelysin-like secreted iron uptake system, extracellular polysaccharide biosynthesis locus |

To better understand the unique biology of the newly identified species, we identified genes likely to be core to novel species but unique within the *Enterococcus* pan-genome were identified in an effort to understand the unique biology of new species. For this, genes sharing <65% amino acid sequence identity over the full-length inferred protein sequence with genes occurring in any enterococcal genome (a cutoff corresponding to the ANI distance between most distant enterococcal species), and which did not occur in proximity to known mobile elements, were identified as likely unique and core to each novel species (Table S6). A total of 1,276 such genes were identified as potentially species-specifying (Table S6), expanding the pan-genome of our sample set of enterococci by 9%. Although many novel genes were of unknown function, we were able to assign a functional annotation to 62% using a combination of annotation and structure-based prediction. This revealed several notable functions encoded by novel species (Table 1, Table S6).

Six new members of Clade I were identified, including candidate species *Enterococcus sp. nov. clewelli* isolated from a captive African Bullet cockroach (Table 1). This species forms an early branch within Clade I in a lineage that gave rise to all Clade I species except for *E. faecalis*. The closest known species to *E. clewelli* is *E. quebecensis*, although the long branch length separating these two species suggests that *E. clewelii* is much more distantly related to *E. quebecensis* than are other novel species and their nearest known relatives. Moreover, only just over half (2234 of 4197) of the predicted genes present in *E. clewelli* occur in *E. rivorum*, highlighting the extensive diversification between these neighboring species. Genes unique to *E. clewelli* encode putative quorum signaling peptides often used in pheromone-responsive plasmids, and the Rnf complex, which is a sodium-coupled ferredoxin oxidoreductase involved in cellular energy generation (Table S6). In addition, the *E. clewelli* genome harbors 233 genes novel to the *Enterococcus* pan-genome, including a cluster of 20 genes which included an iron-dependent peroxidase and proteins associated with adhesion, such as a protein having a predicted von Willebrand factor type A domain and a putative normocyte-binding protein (Table S6).

The second novel Clade I species, candidate species *Enterococcus sp. nov. dunnyi* was isolated twice in our sampling: once from a ground beetle, and once from a captive African Bullet cockroach (Table 1). Its closest taxonomic relative, *E. termitis*, is known to colonize the guts of termites (Švec et al., 2006). *E. dunnyi* contains 603 genes absent from *E. termitis*, representing 20% of its genome content. Gene functions found in *E. dunnyi* but not *E. termitis* include transporters for phosphonate (Table S5, A5889_003100T0-A5889_003103T0) and genes to catabolize dihydroxyacetone (Table S5, A5889_002567T0-A5889_002570T0). Phosphonates are a class of chemicals that include herbicides, and a similar gene cluster confers to *Escherichia coli* the ability to metabolize organophosphates (Jochimsen et al., 2011). *E. dunnyi* contributes 49 genes novel to the *Enterococcus* pan-genome, including two putative glyoxylases potentially involved in resistance to the antimicrobial bleomycin (Table S6).

The third novel Clade I species, candidate *Enterococcus sp. nov. mansoni* was isolated from the droppings of a turkey. Its closest taxonomic relative, *E. quebecensis*, was originally isolated from contaminated water samples (Sistek et al., 2012). The genome of *E. mansoni* harbors 951 genes not shared with *E. quebecensis*, or 30% of its genome, as well as 69 genes novel to the *Enterococcus* pan-genome. Genes present in *E. mansoni* but not *E. quebecensis* include a biosynthetic gene cluster for histidine (Table S5, A5880_000374T0-A5880_000382T0). Genes novel to the pan-genome include a putative toxin similar to the eukaryotic delta conotoxins (Table S6): a class of peptides that targets sodium-gated ion channels (Al-Sabi et al., 2006).

The fourth novel Clade I species candidate *Enterococcus sp. nov. ikei*, most closely related to *E. ureilyticus*, was isolated twice in our collection from a captive moth and a wild dragonfly. Its genome contains 516 genes not shared with *E. ureilyticus* (Table S5), and 14 genes novel to the *Enterococcus* pan-genome (Table S6). The genome of *E. ikei* encodes a putative lactate dehydrogenase complex (Table S5, DIV0869a_001543T0-DIV0869a_001545T0) and catalase (DIV0869a_001489T0). Both lactate dehydrogenase and catalase are important for stress resistance and virulence in *E. faecalis* (Baureder and Hederstedt, 2012; Rana et al., 2013). Pan-genome novel genes include a putative *Listeria*-like autolysin, as well as a miraculin-like protein (Table S6). Miraculins are taste-modifying proteins found in plants (Misaka, 2013), to our knowledge not previously described in bacteria.

The fifth novel Clade I species candidate, *Enterococcus sp. nov. lemimoi*, isolated from a moth and two dragonflies, was found to be a very close taxonomic relative of *E. rotai*. *E. rotai* has previously been isolated from mosquitos, water, and plants (Sedláček et al., 2013). The genome of *E. lemimoi* differs from *E. rotai* by 435 genes, or 14% of total gene content (Table S5). These include a biosynthetic path from oxaloacetate to the amino acid precursor molecule alpha-ketoglutarate (A5866_001933T0-A5866_001946T0). *E. lemimoi* also harbors 20 genes unique to the *Enterococcus* pan-genome (Table S6), including 2 that encoded a putative E1-E2 ATPase, which in many bacteria works as an efflux pumps involved in detoxifying the environment (Silver et al., 1989).

The sixth novel species added to Clade I was candidate *Enterococcus sp. nov. palustris*, isolated from a dragonfly, which branches ancestrally to *E. haemoperoxidus* (Table 1). The gene content of *E. palustris* includes 928 genes not found in *E. haemoperoxidus*: nearly a third of total gene content (Table S5). These species-distinguishing functions include a putative vitamin B12-dependent metabolic pathway to catabolize ethanolamine, as also found in *E. faecalis* (Fox et al., 2009), as well as biosynthetic genes for vitamin B12 (A5821_001318T0-A5821_001343T0). *E. palustris* encodes 33 genes novel to the *Enterococcus* pan-genome, including a putative spherulin-like domain protein (Table S6), a fold that occurs in bacterial proteins associated with stress resilience (Jaenicke and Slingsby, 2001).

Four new members of Clade II were identified. These include candidate *Enterococcus sp. nov. huntleyi*, which was isolated from chicken feces and is most closely related to *E. phoeniculicola* (80.2% ANI; Table 1). The genome of *E. huntleyi*, however, differs from that of *E. phoeniculicola* by more than 50% of its gene content (1573 genes, Table S5), with only 80% identity across shared genes (Table 1). In addition to differences in substrate transport, *E. huntleyi* encodes a metabolic pathway for the catabolism of galacturonate into pentose phosphate pathway precursors (EntecaroJM4C_000098T0- EntecaroJM4C_000100T0) not present in *E. phoeniculicola*, suggesting that this species may be able to metabolize pectin-derived sugars for energy and biomass in a way that its nearest taxonomic neighbor cannot. *E. huntleyi* contributes 84 genes unique to the *Enterococcus* pan-genome (Table S6), including a putative macrolide efflux protein.

The second novel Clade II species, candidate *Enterococcus sp. nov. courvalini*, was isolated from a cockroach and from turkey droppings, and was most closely related to *E. thailandicus*. *E. courvalini* gene content differs from that of *E. thailandicus* by 60 genes (Table S5). These species-differentiating genes encode putative quorum signaling peptides and carbohydrate transporters, functional categories that distinguished many novel species from their nearest neighbors, as well as a putative exfoliative toxin that shares sequence similarity with *Staphylococcus aureus* toxin A/B (DIV0660c_000258T0). *E. courvalini* contributes three new genes to the *Enterococcus* pan-genome, including one encoding a homolog of the restriction endonuclease AseI (Table S6).

The third novel species of Clade II, candidate *Enterococcus sp. nov. wittei*, was isolated from chicken droppings. It was one of two novel species closely related to *E. mundtii*. The *E. wittei* genome contains 783 genes that *E. mundtii* did not (Table S5), representing 30% of the total gene content of this organism. A large gene cluster encoding potassium, nickel and fluoride transport systems, and an acid-activated urea channel and urease (A5844_001283T0- A5844_001298T0), are among those present in *E. wittei* but not *E. mundtii*. This species also harbors 96 genes novel to the pan-genome (Table S6), including three genes encoding different putative protein toxins: a lantarcin-like protein, a thiol activated cytolysin-like protein, and a pheomycin-like protein.

The fourth novel species of Clade II, also closely related to *E. mundtii*, candidate species *Enterococcus sp. nov. mangumii*, was isolated from two butterflies and one dragonfly in our collection (Table 1). It codes for only 321 genes that *E. mundtii* does not, or 11% of its genome (Table S5). These include a gene cluster coding for a metabolic pathway from 2-oxobutanoate, a breakdown product of amino acid catabolism, to 1-propanol (DIV1094d_002603T0- DIV1094d_002624T0). This species additionally contributes 38 genes to the *Enterococcus* pan-genome (Table S6), including a putative discoidin-domain containing protein. Bacterial discoidin domains are associated with lectin binding and adhesion (Cheng et al., 2009).

In Clade III, three new species were identified. Interestingly, all three novel species were isolated from Kemp’s ridley sea turtles that had stranded on the coast of Massachusetts due to cold-stunning. These species included candidate *Enterococcus sp. nov. moelleringi*, which appears to branch early on from the rest of the Clade III strains and is most closely related to *E. pallens*. The gene content of *E. moelleringi* contains 1247 genes absent from *E. pallens*, making up 28% of the novel species genome (Table S5). This gene content includes both aerobic and anaerobic vitamin B12 biosynthetic pathways (Fig. 4B), which were absent from *E. pallens* but present in other Clade III species. In addition to these genes, *E. moelleringi* encodes genes associated with osmoprotection, such as a putative osmolyte transporter (Ent669A_001211T0-Ent669A_001215T0), and glycine/betaine reductase (Ent669A_002128T0- Ent669A_002136T0). In addition, this species encodes a putative V-type ATPase, a family of proteins often used to actively maintain a proton gradient (Ent669A_002615T0-Ent669A_002622T0). *E. moelleringi* contributes gene functions novel to the *Enterococcus* pan-genome, including a collagenase, an acetoacetate decarboxylase, and 130 other genes (Table S6).

The second novel species of Clade III was also most closely related to *E. pallens*. This species, candidate *Enterococcus sp. nov. ferrettii*, branches more recently from *E. pallens*. It encodes 1414 genes absent from *E. pallens*, approximately 28% of its genome (Table S5). These include a nickel/urease gene cluster present in Clade II novel species *E. wittei* (Ent665A_004548T0- Ent665A_004557T0) and a putative fluoroquinolone-specific ABC transporter (Ent665A_004584T0- Ent665A_004586T0). *E. ferrettii* additionally contributes a MAPKKK-like toxin and 108 other genes to the *Enterococcus* pan-genome (Table S6).

The third novel species, candidate *Enterococcus sp. nov. murrayi*, was most closely related to *E. hulanensis*, which is itself a recently discovered species of *Enterococcus* that was isolated from Chinese traditional pickle juice (Li and Gu, 2019). About 22% of the genes in the *E. murrayi* genome (985) are not also present in *E. hulanensis*. Like Clade I novel species *E. dunnyi*, *E. murrayi* encodes homologs of *phnGHIJK*, putatively involved in organophosphonate metabolism (EntMJM16_002940T0- EntMJM16_002951T0 Table S5). *E. murrayi* contributes 80 genes novel to the *Enterococcus* pan-genome (Table S6).

Within Clade IV, we identified five novel species. These include candidate *Enterococcus sp. nov. leclercqi*, isolated from chicken droppings, which is most closely related to *E. diestrammenae*. Genes present in *E. leclercqi* but not *E. diestrammenae* make up about a third of the *E. leclercqi* genome (947 genes) including a polar amino acid transport system (EnteCU9D_000076T0- EnteCU9D_000079T0). The *E. leclercqi* genome also encodes 84 genes novel to the *Enterococcus* pan-genome, including restriction endonuclease Alwl and a putative potential antifungal protein (Table S6).

The second novel species identified in Clade IV, *Enterococcus sp. nov. myersii*, was isolated from aquarium water. It branched ancestrally to species *E. dispar* and *E. canintestini*. Gene functions present in *E. myersii* but not in *E. dispar* or *E. canintestini* represent 21% of the *E. myersii* genome (586 genes Table S5), and include nitrogen regulatory protein and nitrite reductase (EntMJM12_000251T0- EntMJM12_000258T0), a citrate lyase (EntMJM12_000353T0- EntMJM12_000357T0), transport of molybdenum and molybdenum cofactor biosynthesis (EntMJM12_002575T0- EntMJM12_002591T0), and a metabolic pathway from chorismite to indole (EntMJM12_002695T0- EntMJM12_002701T0). The *E. myersii* genome also encodes a novel bacteriocin and 49 other genes new to the *Enterococcus* pan-genome (Table S6).

Candidate *Enterococcus sp. nov. testudinum*, isolated from a captive turtle, was intermediate to *E. casseliflavus /E. flavescens* and *E. gallinarum*. Its closest taxonomic neighbors are the *E. casseliflavus/E. flavescens* group, and *E. testudinum* encodes 713 genes not present in either neighboring species (Table S5). Predicted species-specifying functions in this set include a gene cluster very similar to the molybdenum cofactor biosynthesis genes found in *E. myersii* (A5886_001052T0- A5886_001063T0). *E. testudinum* also encodes a lactococcin-like bacteriocin novel to the *Enterococcus* pan-genome (Table S6).

A fourth novel Clade IV species, candidate *Enterococcus sp. nov. lowellii* also grouped with *E. saccharolyticus* to further define the taxonomic group in Clade IV that branched most recently from the modern-day tetragenococci. As one of the motile enterococci, (Fig. S2) *E. lowellii* was distinct from *E. saccharolyticus* due to the presence of a large flagellar biosynthetic gene cluster (DIV2402_002910T0-DIV2402_002954T0). The genome also encodes a polysaccharide biosynthesis locus novel to the *Enterococcus* pan-genome, suggesting that it may be capable of making a novel type of biofilm or capsule (Table S6).

A fifth Clade IV species, candidate *Enterococcus sp. nov. willemsii*, grouped into a radiation with *E. saccharolyticus* and *E. lowellii*, forming a cluster of species most closely related to the tetragenococci (Fig. 3A). A quarter of the *E. willemsii* genome is not shared with either close taxonomic relative. This functional gene content includes two V-type ATPases (EntCU12B_000279T0- EntCU12B_000289T0 and EntCU12B_000681T0-EntCU12B_000688T0) that are like the species-specifying gene cluster encoded by Clade III species *E. moelleringi*, and a polar amino acid transport system (EntCU12B_002782T0- EntCU12B_002783T0) similar to the species-specifying genes of Clade IV species *E. leclercqi*. *E. willemsii* was found to be one of the motile enterococci (Fig. S2), but the conservation of motility with nearest neighbor *E. lowellii* suggested that the ancestor of both strains may have been motile. *E. willemsii* encodes 58 genes which are novel to the genus (Table S6).

## DISCUSSION

As enterococci occur in the guts of land animals, from mammals to invertebrates (Lebreton et al., 2017; Martin and Mundt, 1972; Mundt, 1963a), host-microbe interactions can be examined and compared in an unusually wide variety of ecological contexts and physiological systems, providing a unique opportunity to discover universal principles that govern host/gut microbe interactions. Further, enterococci have emerged among leading causes of antibiotic resistant hospital associated infection (Fiore et al., 2019; Gaca Anthony O. and Lemos José A., 2019), making understanding of host association principles imperative. Since the work of Mundt and colleagues from the 1960’s and 1970’s, when fewer than 10 enterococcal species had been defined using phenotypic criteria, over 60 species now have been proposed (Parks et al., 2020), with most associated with at least one representative genome sequence. With no exception, known enterococcal species differ from their closest sister species by > 5% average nucleotide identity, the threshold proposed as a new “gold standard” for species definition (Jain et al., 2018). Here we described the isolation and distribution of many known species from diverse hosts and geographies, and further identified 18 new lineages that qualify as new species by the above criterion (Table 1).

The isolation of 18 novel species from diverse wild samples suggests that much species-level diversity within the genus *Enterococcus* remains to be discovered. We used our isolation data, and published estimates of global animal biomass and animal species diversity (Bar-On et al., 2018; Barrowclough et al., 2016; Stork, 2018), to understand the depth to which we had sampled the diversity of different animal taxa, and to extrapolate an upper bound for enterococcal species diversity (Fig. 5A). Wild birds and mammals represent only 3% and 1% of the biomass of terrestrial arthropods respectively, and we recovered fewer different species of *Enterococcus* from those animal groups than from the comparatively small sampling of terrestrial arthropods. A comparison of *Enterococcus* species diversity to the fraction of animal group diversity sampled highlights the depth to which each group was sampled in this study (Fig. 5B). This analysis revealed that mammalian diversity has been generally well sampled with few new enterococcal species arising, but that the tremendous diversity of terrestrial arthropod hosts was particularly fertile with only a hundred-thousandth of a percent of total arthropod host species diversity having been explored. There are only ∼10^3^ existing species of mammals, all with distal gut physiologies that are broadly similar. In stark contrast, there are estimated to be nearly 10^7^ species of arthropods with widely varying diets (carnivorous to highly specialized herbivores) and gut physiologies (ranging from alkaline to acidic). This study found most of the new species in the comparatively few insect and invertebrate samples examined (enterococci were isolated from 140 total terrestrial Arthropod samples, see Table S1), and in insectivorous agricultural poultry that are likely capable of serving as aggregators of insect enterococcal species. Based on the small sample size (140) and limited diversity of insects and invertebrate samples examined here, we would predict the existence of potentially thousands of undiscovered species of enterococci defined as varying from the closest extant relative by >5% ANI.

**Figure 5.**
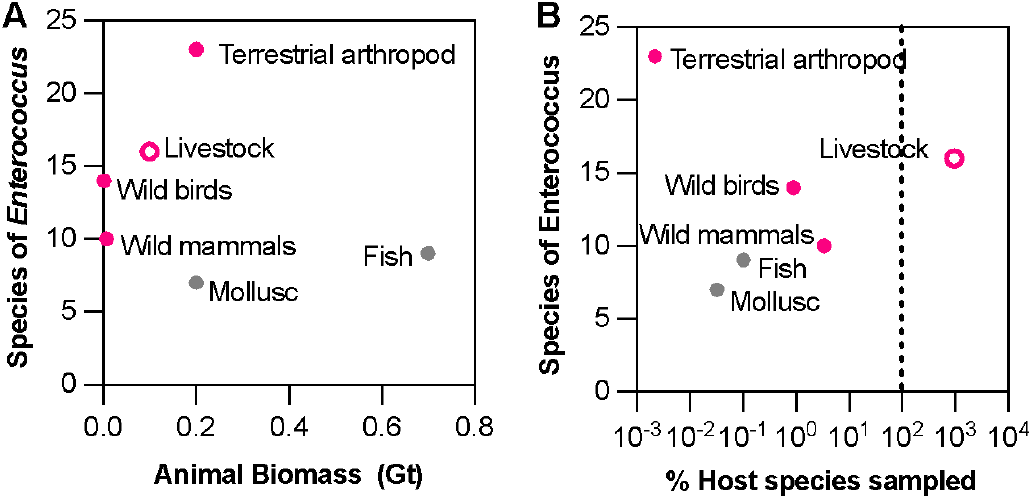
Estimate of enterococcal species abundance and distribution across animal hosts. **A**) Number of *Enterococcus* species identified in each host type, plotted as a function of total global animal biomass that host type represents. B) The number of *Enterococcus* species identified as a function of the fraction of species within each category sampled in this study, showing that terrestrial livestock were over sampled relative to their diversity, whereas the diversity of terrestrial arthropod species was comparatively superficially explored. Dashed vertical line indicates 100%. In both plots: pink filled circle, terrestrial; pink open circle, terrestrial-agricultural; grey circle, aquatic.

In this work, as did Mundt and colleagues before us (Martin and Mundt, 1972; Mundt, 1963a, 1963b, 1961; Mundt et al., 1958), we observed several enterococcal species (e.g., *E. faecalis*, *E. mundti*, and *E. faecium*) associated with a wide variety of hosts, whereas others were rarer, potentially as a consequence of being host specific (Fig. 2A). From the wide distribution and abundance of *E. faecalis*, we deduce that this species is likely a generalist able to proliferate within diverse hosts to large population sizes and survive outside of the host: two factors that allow successful transmission. In contrast, the plethora of rarer species suggests that many enterococci are specialists and are less able to proliferate in diverse host environments, less robust to the stresses associated with transmission outside of a host, or possess traits that allow colonization of specialized host environments. The paths through which generalist and specialist species evolve remain unknown, but our work raises the possibility that host trophic or social interactions gave rise to the observed pattern of generalist and specialist species through the creation of barriers to gene flow.

Hosts themselves vary widely in which *Enterococcus* species they harbor: In contrast to known specific associations, such as that of *E. columbae* with the Columbidae family of pigeons and doves as previously discussed (Lebreton et al., 2017), humans are typically colonized by either *E. faecalis* or *E. faecium*, or both (Dubin Krista et al., 2017). In contrast, chicken and poultry are colonized by a wide diversity of species, as observed here and previously (Rehman et al., 2018). In the present study in addition to 10 known species, we identified 6 new species of *Enterococcus* associated with poultry (Table 1). It seems unlikely that these 6 new species (and as many more established species of *Enterococcus* observed by us and others) each evolved specifically to colonize the poultry gut. In nature, poultry and closely ancestrally related birds are highly adapted foragers that exhibit an innate ground scratching behavior, unearthing insects, and other sources of nutrition (Klasing, 2005). It is plausible that early exposure to microbes in the diet shapes the assembly of gut communities in this context, and that opportunities exist for the transient occurrence of specialist and generalist enterococci to be isolated from poultry feces. A similar phenomenon may also underlie the diversity of enterococcal species that we observed in dragonflies (Table S1). We isolated 10 *Enterococcus* species, including 5 novel species, from this carnivorous order of insect. Dragonflies eat other insects, and the microbiota of their prey has been found to shape their gut microbiota composition (Deb et al., 2019). Together, these results suggest that animals whose diet contains a large proportion of insects and other arthropods may be aggregators and reservoirs of particularly diverse enterococci.

The acquisition of adaptive genes can facilitate genetic differentiation by allowing individuals carrying these genes to proliferate in new ecological niches (Polz et al., 2013). Our finding that many Clade III strains have acquired vitamin B12 biosynthesis pathways suggests that the ability to make this vitamin may be a key trait for the ecology that Clade III strains inhabit (Figure 4B). As we showed, these pathways in fact relieve the B12 auxotrophy that is otherwise common to all other enterococci (Fig. 4C). B12 is rare in diets derived purely from plant or fungal matter (Roth et al., 1996). Cobalamin is made and shared among microbes in various complex consortia (Degnan et al., 2014) including those of the gut. All 3 novel and phylogenetically separate Clade III species newly identified in this study derived from cold-stunned Kemp’s ridley sea turtles that came ashore on Cape Cod. These turtles often live in shallow coastal marine water, and consume a diet of mainly crustaceans, and to a lesser extent plants (Seney and Musick, 2005). The occurrence of 3 separate Clade III species in samples of the same host type from similar locations suggests that they may derive from the diverse coastal crustaceans consumed. An alternative explanation would be that these species are each independently adapted to the Kemp’s ridley turtle, however this seems less likely in a shared ecosystem where one host-adapted species would likely predominate over time. Many crustaceans subsist on a diet of algae, photosynthetic eukaryotes at the base of the marine food chain that are incapable of synthesizing cobalamin (Roth et al., 1996). A cobalamin-synthesizing *Enterococcus* in the guts of coastal algae consumers would seem well placed. Together with the intriguingly large genomes of species within this clade, these data suggest that Clade III enterococci occupy a novel ecological niche compared to other species within the genus.

The prevalence of amino acid auxotrophy including branched chain amino acids and histidine (Figure 4E) is a notable signature of clade II enterococci. For members of the closely related genus *Lactococcus*, auxotrophy for branched chain amino acids and histidine, which are abundant in milk protein, is the hallmark of mammalian-adaptation (Delorme et al., 1993; Godon et al., 1993), a finding that has been recapitulated under laboratory evolution conditions (Bachmann et al., 2012). Our previously proposed timeline for *Enterococcus* evolution positions the radiation that split Clade II from Clades III and IV about 350 million years ago (Lebreton et al., 2017), which would predate the evolution of mammals from synapsid ancestors (approximately 200 million years ago (Álvarez-Carretero et al., 2022)). This suggests that the radiation of Clade II may have resulted from an earlier adaptation by enterococci to the guts of the synapsid precursors of mammals, animals that dominated terrestrial life beginning approximately 318 million years ago (Brocklehurst, 2021). The disappearance of the last surviving non-mammalian synapsid about 30 million years ago (Matsuoka et al., 2016) now leaves only mammalian representatives of that division – and their gut microbes – to be studied, potentially explaining why surviving Clade II enterococcal species share amino acid auxotrophies characteristic of mammalian-adapted lactococci. Of clinical relevance, diets that include lactose have been observed to predispose hospitalized patients to *E. faecium* overgrowth and subsequent infection (Stein-Thoeringer et al., 2019). It is notable that *E. faecalis*, a Clade I species, showed similar patterns of amino acid auxotrophy. Future work will be needed to determine ecological and evolutionary factors that led to these convergent patterns of auxotrophy in the two species of *Enterococcus* in which clinically relevant multidrug resistance has emerged.

Finally, we identified important genetic novelty in widely distributed generalist species of enterococci proximal to the human ecology, *E. faecalis, E. faecium and E. hirae*, with implications for human health. From strains collected for this study we separately identified an entirely new class of botulinum-type toxin encoded on a plasmid that occurred in an *E. faecium* excreted from the gut of a cow (Zhang et al., 2018). We separately identified a new class of pore forming toxins distantly related to the delta toxins of clostridia, in domesticated animal-derived strains of *E. faecalis, E. faecium and E. hirae* (Xiong et al., 2022), the three most commonly encountered generalists in this study. The role of either of these novel classes of toxins in the natural biology of enterococci remains to be discovered, but the discovery of potential biological threats in such ubiquitous and easily transmitted gut microbes associated with human infection is disconcerting. This is especially true given that the implied genetic diversity is vast in the large planetary burden of a microbe that is a ubiquitous component of diverse host-associated microbiome populations. Expanding this search now in a more informed and directed way will illuminate the scope of enterococcal genetic diversity and potential sources and routes of transmission of antibiotic resistance genes, providing new insight into the mechanistic basis for host association of this important class of microbes.

## METHODS

### Isolation of a library of diverse enterococci

Approximately one gram of fecal sample, small whole insects, or gut tracts dissected from large invertebrate samples were mechanically disrupted using a mortar and pestle in 1-2 volumes of sterile pH 7.4 phosphate buffered saline. Sample dissections were carried out with sterilized equipment. Glycerol was added to homogenate as a cryoprotectant, and remaining material was stored at −80 C. Dilutions of the homogenate were directly plated onto bile esculin agar (BEA; Difco 299068). This medium contains oxgall bile as a selective agent and produces a differential colorimetric reaction upon esculin hydrolysis. Bile resistance is a trait that defines the genus *Enterococcus* (Fiore et al., 2019), and many species hydrolyze esculin. Enterococci were enriched by plating a 100 µl aliquot of the homogenate onto a 25 mm #1 Whatman filter paper placed on a BEA agar plate. The filter papers were incubated at room temperature for 48-72 hours, or until black pigment developed. Bacterial growth was collected by rinsing filters in brain-heart infusion (BHI) medium (Difco DF0418) and plated onto BEA agar to retrieve single colonies. Morphologically distinct isolated colonies were selected and purified by a further plating onto BEA or CHROMagar Orientation medium. Isolates that grew on BEA, or that produced blue pigmented colonies on CHROMagar were selected for taxonomic identification. Isolates were routinely maintained in BHI, and frozen at −80 °C in BHI containing 20% glycerol for long-term storage.

### Identification of isolates

As a template for molecular analysis, genomic DNA was collected from cells grown in BHI with a Blood and Tissue kit (Qiagen), following the manufacturer supplied protocol with the addition of mutanolysin to fully disrupt enterococcal cell walls during cell lysis. Isolates positively identified as *Enterococcus* spp. were selected for further analysis. Orthologs of *E. faecalis* V583 gene EF1948 present in a sample set of 47 species of *Enterococcus* were aligned to identify conserved regions, and oligonucleotide primers were designed to amplify a conserved region flanking 97 bp of more variable sequence called diversity locus (DL; Table S7). The annealing temperature of the primers was optimized by gradient PCR using genomic DNA from 24 known enterococcal species, as well as DNA from 8 non-enterococcal controls from the genera *Streptococcus, Lactococcus*, *Vagococcus* and *Carnobacteria*. An annealing temperature of 51 °C correctly amplified a single product from each enterococcal template but did not amplify products from the non-enterococcal controls. Where the DL primers failed to amplify a product, the full-length 16S rDNA sequence amplified with primers 8F and 1522R (Turner et al., 1999) with High Fidelity Q5 Taq polymerase (NEB). A high-quality sequence was obtained by the Sanger method and was used to query the 16S rRNA BLAST database. In all instances, isolates that failed to amplify a product were found to be species other than *Enterococcus*.

### Genome sequencing and assembly

For 33 isolates, genomic DNA was used to generate dual indexed Nextera-XT short-read sequencing libraries (Illumina). We sequenced these libraries in paired end format with 250 bp read length. We performed de novo assembly of short reads using CLC Workbench, after trimming adapter sequences, filtering reads based on quality score and removing PhiX. For the remaining 14 isolates, we prepared whole-genome paired-end and mate-pair Illumina libraries, and sequenced and assembled them as previously described (Lebreton et al., 2017). Sequencing reads and assemblies were submitted to GenBank under BioProjects PRJNA324269 and PRJNA313452.

### *In vitro* growth assays for *Enterococcus* spp. on medium with or without cobalamin

A total of 4 enterococcal strains (*E. faecalis* V583, *E. gilvus* BAA-350, *E. avium* ATCC14025 and *E. raffinosus* ATCC49464) were grown on BUG+B agar plates at 30°C. Cells were harvested and resuspended in 5 ml PBS to reach OD590 = 0.1. Cell suspensions were centrifuged, washed twice, and resuspended in 1:1 PBS. A 1% inoculum from each cell suspension was inoculated in a vitamin B12-free chemically defined medium (CDM) and grown overnight at 37°C. The vitamin B12-free medium was modified from an existing CDM used for lactic acid bacteria (PMID: 21097579) and contained the following per liter: 1 g K_2_HPO_4_, 5 g KH_2_PO_4_, 0.6 g ammonium citrate, 1 g acetate, 0.25 g tyrosine, 0.24 g alanine, 0.125 g arginine, 0.42 g aspartic acid, 0.13 g cysteine, 0.5 g glutamic acid, 0.15 g histidine, 0.21 g isoleucine, 0.475 g leucine, 0.44 g lysine, 0.275 phenylalanine, 0.675 g proline, 0.34 g serine, 0.225 g threonine, 0.05 g tryptophan, 0.325 g valine, 0.175 g glycine, 0.125 g methionine, 0.1 g asparagine, 0.2 g glutamine, 10 g glucose, 0.5 g l-ascorbic acid, 35 mg adenine sulfate, 27 mg guanine, 22 mg uracil, 50 mg cystine, 50 mg xanthine, 2.5 mg d-biotin, 1 mg riboflavin, 5 mg pyridoxamine-HCl, 10 μg *p*-aminobenzoic acid, 1 mg pantothenate, 5 mg inosine, 1 mg nicotinic acid, 5 mg orotic acid, 2 mg pyridoxine, 1 mg thiamine, 2.5 mg lipoic acid, 5 mg thymidine, 200 mg MgCl_2_, 50 mg CaCl_2_, 16 mg MnCl_2_, 3 mg FeCl_3_, 5 mg FeCl_2_, 5 mg ZnSO_4_, 2.5 mg CoSO_4_, and 2.5 mg CuSO_4_.

When necessary, vitamin B12 was supplemented with the addition of 100, 10 or 1 μg cyanocobalamin per 100 mL (CN-Cbl, Sigma USA). Optical density at 590 nm (OD590) was monitored hourly for 24 h, using a Synergy 2 microplate reader (Bio-Tek).

### Comparative Genomics

#### Selection of comparator genomes

We selected known species of *Enterococcus* and other members of the taxonomic family Enterococcaceae that were also found in guts, choosing those with assemblies of at least 96% completeness, as estimated by CheckM (Parks et al., 2015). In many cases, a single representative meeting this criterion was available for each species. Where multiple representatives of a species were sequenced, we selected a representative for each. However, because *E. casseliflavus* genomes showed a high degree of intra-specific variation in ANI, we selected assemblies from three strains of *E. casseliflavus* isolated from human, plant, and chicken sources (Table S2), as well as an assembly of *E. flavescens*, which has previously been proposed as a species of *Enterococcus* but is very closely related to *E. casseliflavus* (Naser et al., 2006; Pompei et al., 1992). We added a total of 33 comparator genomes, resulting in a total of 103 genomes for comparative analysis.

#### Annotation of genomes

We uniformly annotated the 47 newly sequenced *Enterococcus* and *Vagococcus* isolates, as well as the set of comparator genomes, using the Broad Institute’s prokaryotic annotation pipeline (Lebreton et al., 2013). Furthermore, we annotated coding regions using i) KoalaBLAST to identify genes with KEGG annotations (Kanehisa et al., 2016; Kanehisa and Goto, 2000); ii) the CARD database to annotate antibiotic resistance genes using the RGI tool (Jia et al., 2017); iii) dbCAN to identify carbohydrate active enzymes described in the the CAZy database (Cantarel et al., 2009; Zhang et al., 2018); iv) AntiSMASH to identify secondary metabolite gene clusters, including lanthipeptide synthases (Medema et al., 2011); v) CRISPRDetect to identify CRISPR spacers (Biswas et al., 2016); and vi) ProphET to identify prophage (Reis-Cunha et al., 2019).

#### Calculation of average nucleotide identity

To calculate average nucleotide identity (ANI) between the whole genome sequences in our dataset, we used FastANI (Jain et al., 2018), which performs a kmer-based comparison.

#### Clustering of orthologous genes and core gene phylogeny

We identified orthologous clusters of genes across our complete set of 103 genomes using OrthoFinder (Emms and Kelly, 2015), which is optimized for highly diverse datasets. In order to generate a phylogenetic tree, we identified the set of 320 orthologous groups representing genes found in single copy across all isolates (i.e., single copy core; SCC), performed multiple-sequence alignment using MAFFT-linsi (Katoh and Standley, 2013), converted this alignment to a codon-based alignment using PAL2NAL (Suyama et al., 2006), and then used this alignment to construct a phylogenetic tree using IQ-TREE (beta-1.7) (Minh et al., 2020) with 1000 bootstrap replicates, using ModelFinder Plus to find the best codon model. Phylogenies were visualized using iTOL (Letunic and Bork, 2019, 2016).

#### Clade-level enrichment analysis

We identified traits enriched or depleted in specific enterococcal clades, as well as differentiating the entire genus from outgroups by requiring a trait to be present in at least 80% of the members of one clade and at most 20% of the members of the comparing clade. Traits tested for differential presence between taxonomic clades included: KEGG Ortholog Groups, KEGG Modules, KEGG Pathways, OrthoFinder Ortholog Groups, CARD AMR alleles, and CAZy Carbohydrate Utilization Enzymes.

#### Comparison of novel species gene content to nearest taxonomic neighbor

The shared gene content of comparator species was calculated by reciprocal BLAST of gene sequences, using a cutoff of 64% identity across full-length nucleotide sequences of annotated genes. This % cutoff was used because the two least related enterococci were found to share 65% average shared nucleotide identity. Genes identified by reciprocal BLAST were identified as being present in both genomes. Where multiple comparator genomes existed, genes had to be present in all comparators to be considered shared, or present only in novel species comparators to be considered part of the set that differentiated the novel species from its nearest taxonomic neighbor. The predicted function of genes present only in novel species was further analyzed by mapping COG annotations to their letter code, which defined larger functional classes.

#### Analysis of gene content characterizing novel species

To investigate the unique characteristics encoded in the novel species, we identified and characterized 1426 genes that were found in a novel species, and in no other enterococcal species, using our orthogroup clusters. In addition to the functional annotation described in “Annotation of genomes” we performed structure-based function prediction for genes of unknown function using PHYRE2 with a confidence or sequence identity threshold of 90% (Kelley et al., 2015). To identify genes which may have originated from horizontal gene transfer, we searched for mobile elements, including Prophages (see Annotation of genomes section above) and IS elements in close proximity (10 genes upstream or downstream) using ISfinder (Siguier et al., 2006). We also identified clusters of these genes which colocalized on the chromosome of a particular novel species, or genes which were at most 10 genes apart on the same scaffold.

#### Estimating species diversity within the genus *Enterococcus*

We gathered estimates of total animal biomass, and species diversity within taxonomic groups of animals from the literature (Bar-On et al., 2018; Barrowclough et al., 2016; Burgin et al., 2018; Stork, 2018). We estimated the depth to which we had sampled animal species diversity by dividing the total number of samples collected for one animal type by the published estimates of total species. This approximation overestimates the depth of sampling where we sampled multiple animals belonging to the same species. We chose this method because it was not possible to make a species-level taxonomic identification for some sample types, such as the scat of wild animals or some whole arthropod samples.

## ACKNOWLEDGEMENTS

Samples were collected by the Enterococcal Diversity Consortium, a worldwide consortium of collaborators, including Adventure Scientists, Consumer Reports, the Marine Resources Center, the New England Aquarium, the Clemson University Morgan Poultry Center, the Eisen Laboratory, the Kolter Laboratory, Alexander Bertonneau, Kristal Bertonneau, Sophia Bertonneau, Peter Billman, Allen Bolinger, Robert Bruker, Ilana Camargo, Peter Claussen, Tucker Cunningham, Lonnie Dupre, Colleen Ferris, Nkrumah Frazier, Ana Frazzon, Matt Gaidica, Marla Garrison, Rebecca Gast, Michael Gilmore, Peter Girguis, Gonzalo Giribet, Erin Gontang, Jennie Groves, Sebastian Gunther, Wolfgang Haas, Devin Huntley, Suzie Imber, Charles Innis, Mike Libecki, Abigail Manson, Joseph Manson, Pascale Marceau, Megan May, Nathan McGuire, Patrick McGuire, Richard McLaughlin, Philip Metzger, Joanne Munisteri, Andrew Oster, Janira Prichula, Matthew Rowbottom, Stefani Ryan, Katharina Schaufler, Róza Sebök, Helen Seneker, Bruna Sgardioli, Ken Tennessen, Gregg Treinish, Daria Van Tyne, Hera Vlamakis, Jeff Vohl, Jaap Wagenaar, Maarten Gilbert, Jenna Wallenga, Jeremy Wei, and Sheila Withrow. Those enterococci isolated in Sao Carlos, Brazil are registered at the National System for the Management of Genetic Heritage and Associated Traditional Knowledge of Brazil, SISGEN, under the number A85F977, and those isolated in Porto Alegre, Brazil under SISGEN number A720680.

This project was funded by the Harvard-wide Program on Antibiotic Resistance NIH/NIAID grant AI083214, and U19AI110818 to the Broad Institute. JAS was supported by the National Institutes of Health Ruth Kirschstein fellowship F32GM121005.

