## Supplemental Figures 1-3 for "Global diversity of enterococci and description of 18 novel species"

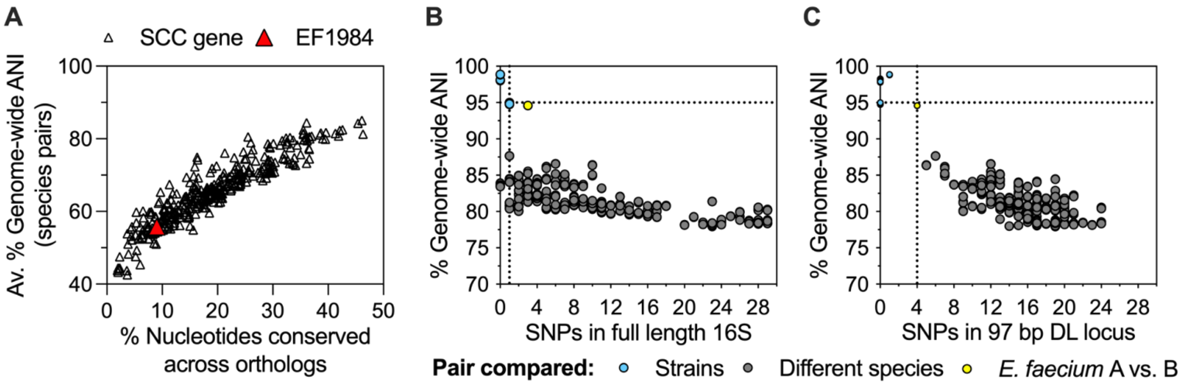


**Figure S1. Identification of a molecular marker for species-level identification of *Enterococcus*. A**) Selected *Enterococcus*-specific marker gene, EF1984 (red triangle), compared to 1037 genes shared in single copy by 28 species of *Enterococcus* (open triangles, (Lebreton et al. 2017)). The percent of identical nucleotides shared by a multiple alignment of each gene is plotted against the average pairwise nucleotide sequence to highlight that, compared to other genes core to *Enterococcus*, the EF1984 sequence is highly variable: pairs of species share on average 55% nucleotide identity. Within EF1984 we identified a 97 bp Diversity Locus (DL) sufficient to screen isolates for diverse species of *Enterococcus.* **B-C**) Using a previously sequenced collection of 42 diverse enterococcal isolates, these plots show the ability of single nucleotide polymorphism (SNP) counts within either the full-length 16S rRNA gene (B) or the 97 bp DL locus C) to predict the genome-wide average nucleotide identity (ANI). Same and different-species relationships among pairs of strains of the same (gray) or different (other color) species are indicated. The species threshold based on ANI (95%) is shown with a dashed horizontal line. Dashed vertical lines indicate 1 SNP in full-length 16S (B) or 4 SNP in DL (C). SNPs are negatively correlated with genome wide average nucleotide identity. (Pearson correlation: DL, r(282)= -0.63, p=<0.0001; 16S, r(373)=-0.59, p=<0.0001)

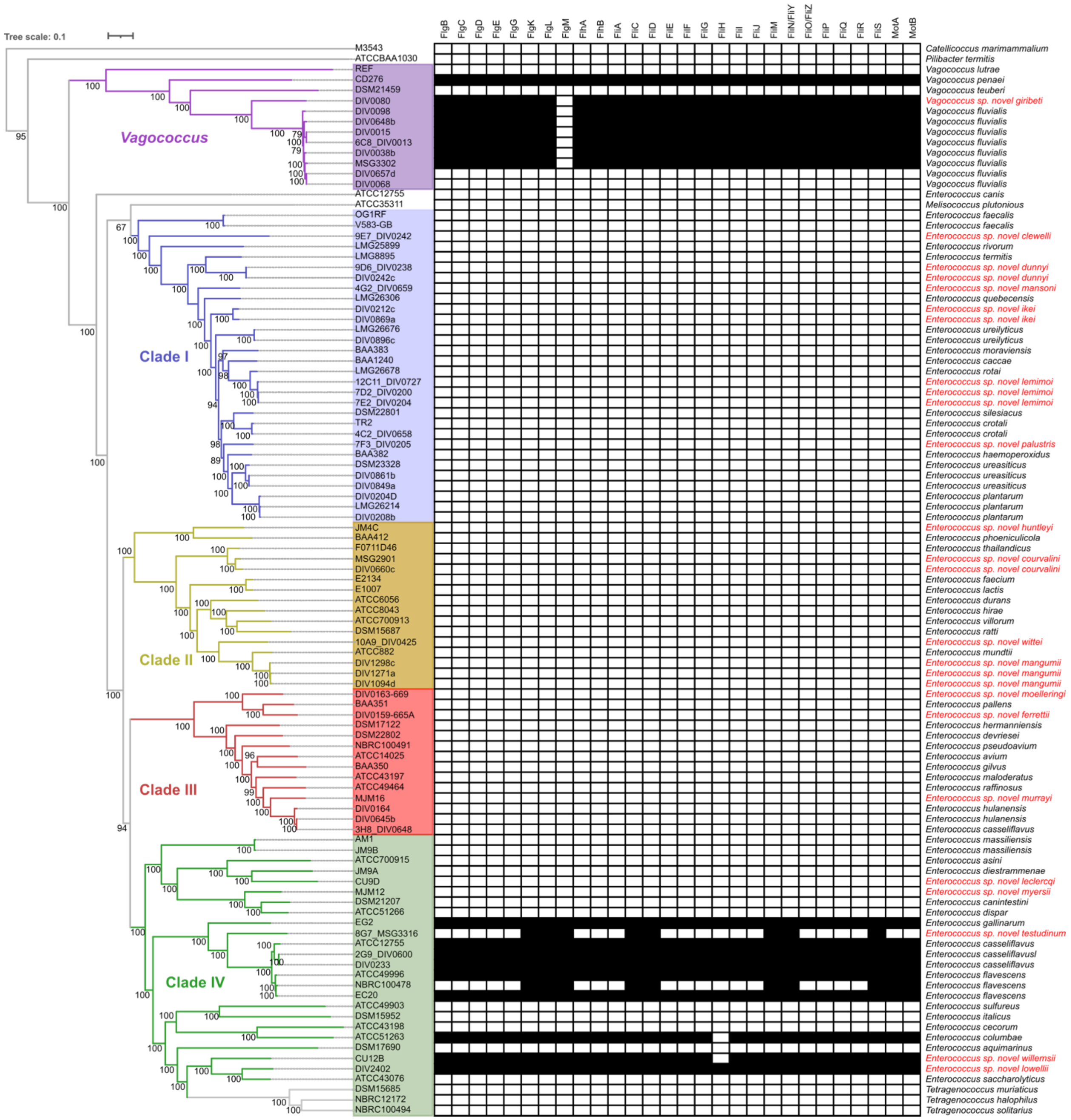

**Figure S2. Conservation of flagellar motility orthologous gene content between Vagococcus and clade IV *Enterococcus* species.** Concatenated core gene phylogeny of *Enterococcus*, as shown in Figure 3. Red text indicates novel species. Numbers annotating phylogeny indicate bootstrap support from 1000 replicates. Conservation of orthologous genes related to the bacterial flagellum and motor are shown as filled black boxes.

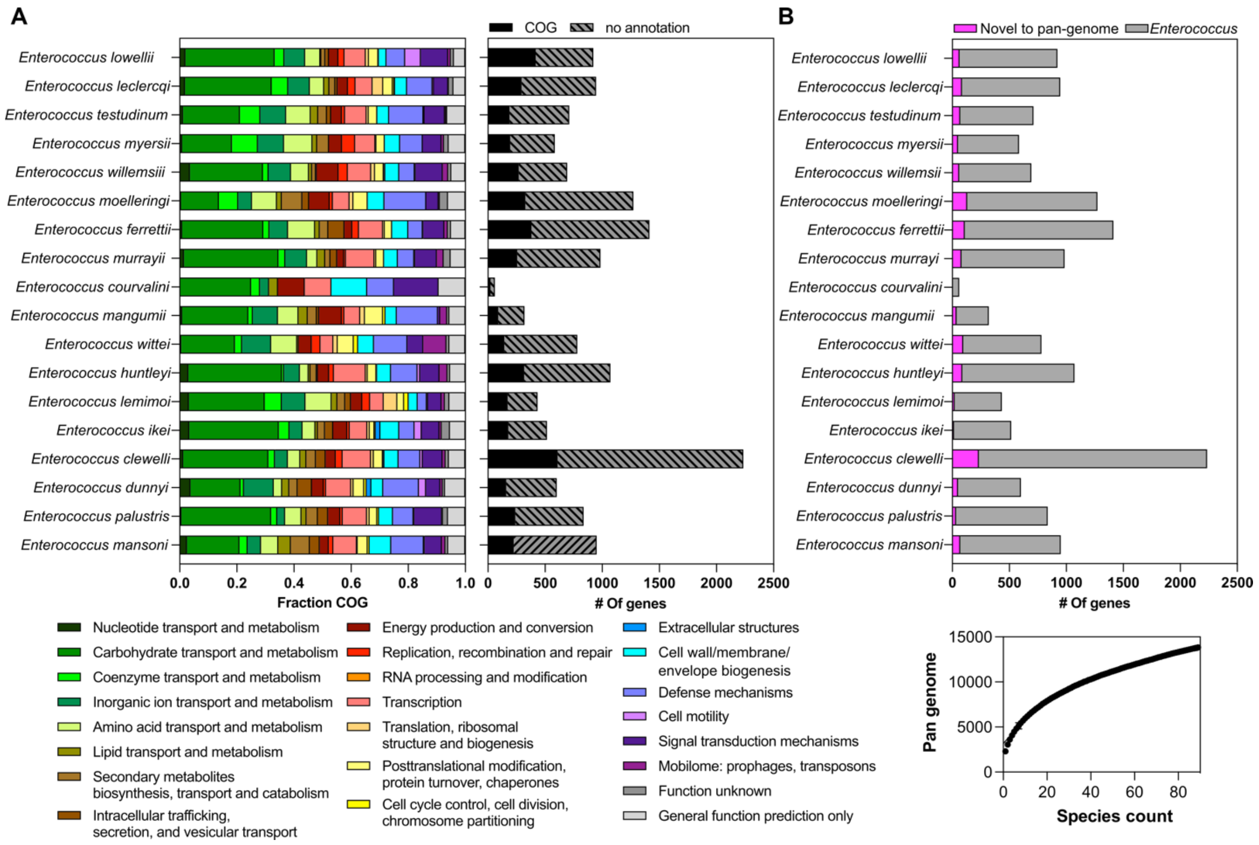


**Figure S3. Enrichment of functional gene categories and expansion of the pan genome by novel *Enterococcus* species.** A) Gene content encoded by a novel species but not its nearest taxonomic neighbors, binned by functional category as defined by COG (left). The total number of 'species specifying' genes, and the proportion of these genes with a COG annotation is shown at right. B) Species-specifying gene content that contributes novel orthologous gene groups to the *Enterococcus* pan genome. Rarefaction curve shows the gene-level diversity captured by the phylogeny presented in figure 3. Genes novel to the *Enterococcus* pan genome are defined as the set that is present only in the genomes of novel species and absent from the genome of any other Enterococcus species deposited in ncbi.
