## Supplementary material for "Global diversity of enterococci and description of 18 novel species": Table S7

| Primer | Bases | Tm |
| --- | --- | --- |
| EF3480F1 | GGTGTGGATATTGATGAAAC | 51 |
| EF3480F2 | GGTGTGGATAATGATGAGAC | 51 |
| EF3480F3 | GGTGTGGATATTGACGAAAC | 51 |
| EF3480F4 | GGTGTGGATATTGATGATAC | 51 |
| EF3480F5 | GGCGTGGATAACGACGAAAC | 51 |
| EF3480F6 | GGTGTTGACATTGATGAAAC | 51 |
| EF3480R1 | TCATCATTTGGATAATAGCC | 51 |
| EF3480R2 | TCATCTTGTGGATAATAGCC | 51 |
| EF3480R3 | TGATCATTTGGATAATACCC | 51 |
| EF3480R4 | TGATCATTCGGATAATAACC | 51 |
| EF3480R5 | TCATCATTCGGGTAATACCC | 51 |

Primers used to amplify DL locus for molecular typing of *Enterococcus* species.
